# A systems genetics approach identifies roles for proteasome factors in heart development and congenital heart defects

**DOI:** 10.1101/2025.01.16.633339

**Authors:** Gist H. Farr, Whitaker Reid, Isabelle Young, Mona L. Li, David R. Beier, Lisa Maves

## Abstract

Congenital heart defects (CHDs) occur in about 1% of live births and are the leading cause of infant death due to birth defects. While there have been remarkable efforts to pursue large-scale whole-exome and genome sequencing studies on CHD patient cohorts, it is estimated that these approaches have thus far accounted for only about 50% of the genetic contribution to CHDs. We sought to take a new approach to identify genetic causes of CHDs. By combining analyses of genes that are under strong selective constraint along with published embryonic heart transcriptomes, we identified over 200 new candidate genes for CHDs. We utilized protein-protein interaction (PPI) network analysis to identify a functionally-related subnetwork consisting of known CHD genes as well as genes encoding proteasome factors, in particular *POMP*, *PSMA6*, *PSMA7*, *PSMD3*, and *PSMD6*. We used CRISPR screening in zebrafish embryos to preliminarily identify roles for the PPI subnetwork genes in heart development. We then used CRISPR to create new mutant zebrafish strains for two of the proteasome genes in the subnetwork: *pomp* and *psmd6*. Phenotypic analyses confirm critical roles for *pomp* and *psmd6* in heart development. In particular, we find defects in myocardial cell shapes and in outflow tract development in *pomp* and *psmd6* mutant zebrafish embryos, and these phenotypes have been observed in other zebrafish CHD-gene mutants. Our study provides a novel systems genetics approach to further our understanding of the genetic causes of human CHDs.

**Author Summary:** Congenital heart defects (CHDs) are birth defects resulting in the abnormal structure and function of the heart. Genetic mutations are a significant cause of CHDs. Many studies have used genome sequencing of CHD patients and their families to gain knowledge of the mutations that cause CHDs. However, these studies have only found about 50 percent of the genes involved in CHDs. Here, we take a new approach to identifying genes that are required for heart development and that may cause CHDs, generating a list of over 200 candidate genes. Using multiple data systems, including human exome sequences, mouse transcriptomes, and protein-protein interactions, we identify a small group of related potential CHD genes that includes multiple genes encoding proteasome factors. These factors are known to be important for assembling the proteasome, a large molecular machine that eliminates unneeded or damaged proteins from the cell, but which has not been shown to contribute to CHD. We use a CRISPR-based approach in zebrafish to specifically eliminate some of these candidate genes and reveal new roles for proteasome genes in heart development. We show that loss of proteasome gene functions leads to zebrafish heart defects that resemble those seen in other zebrafish CHD-gene mutants. This study shows that a proteasome gene family contributes to heart development, advancing our understanding of the causes of CHDs. By increasing our understanding of the genetic causes of CHDs, our work should lead to better screening, more accurate diagnoses, and, ultimately, better treatments for these disorders.

## Introduction

Congenital heart defects (CHDs) occur in about 1% of live births and are the leading cause of infant death due to birth defects [1–4]. CHDs are structural malformations of the heart that result from disruptions in cardiac development. There is a broad spectrum of CHDs, ranging from those affecting a particular valve or chamber to more severe and complex abnormalities involving multiple heart chambers and vessels [5]. CHDs may occur as isolated malformations or in combination with extracardiac defects [5–7]. Although environmental causes can contribute to CHDs, numerous studies point to a strong genetic component and high heritability for many forms of CHDs [8–12]. Identifying the genetic causes of CHD can not only provide better understanding of heart development but also can inform genetic counseling and clinical care.

Several types of genetic alterations have been shown to contribute to CHDs, including chromosomal aneuploidies, copy number variants, small insertions or deletions, as well as *de novo* and inherited single-nucleotide variants [7,12,13]. Large-scale whole-exome sequencing studies find that CHD cases show an excess of damaging coding *de novo* variants (DNVs) as well as rare inherited loss-of-function coding mutations [14–19]. These studies estimate that about 10% of CHD cases may be caused by coding DNVs, while whole genome sequencing studies estimate that noncoding DNVs may also confer a substantial contribution to CHDs [20,21]. Taken together, it is estimated that these CHD patient cohort sequencing studies have thus far accounted for about 50% of the genetic contribution to CHDs [12].

Many approaches have been taken to define the “known” CHD genes [13]. One study has defined high-confidence CHD genes as genes in which variants have been reported as the monogenic cause of CHD in at least 3 independent familial or sporadic cases, in one or more separate publications (132 genes at the time of this study; [22]). However, it is estimated that there are over 440 risk genes for CHDs [18]. Therefore, a major hurdle that remains for understanding the causes of CHDs is the identification and validation of human CHD genes that are as yet unknown.

As a complementary approach to sequencing patient cohorts for human disease-gene discovery, larger-scale human population genome data has been analyzed for essential genes [23–26]. Such studies find that genes that are essential for mammalian embryonic development are strongly correlated with human disease genes, especially for developmental disorders [25]. An analysis of the ExAC human exome sequencing database identified a set of genes for which heterozygosity for nonsense mutations is rare or absent in the normal adult population [23].

These genes are thus predicted to be haploinsufficient in humans and are strong candidates for contributing to birth defects. From this analysis, Cassa et al. assigned a heterozygote selection value (*s*_het_ score) to each human gene in the genome [23]. The *s*_het_ score correlates with known human autosomal dominant disorders, indicating the relevance of genes with high heterozygote selection values to human disease [23]. The *s*_het_ score also correlates with developmental lethality in homozygous mouse knockout strains generated by the International Mouse Phenotyping Consortium (IMPC) [23,27]. Thus, these analyses implicate high *s*_het_ genes in mammalian development and disease.

As a complement to mouse models, zebrafish offer many advantages for screening and characterizing new CHD genes. Zebrafish provide the ability to examine the earliest stages of heart development in live externally-developing embryos, and many studies have documented that genes needed for proper heart development in zebrafish also contribute to CHDs in humans [28–30]. Another major advantage of the zebrafish model is the availability of efficient CRISPR screening and mutagenesis approaches, which have shown success for candidate CHD gene discovery [29,31,32].

Here we take a novel approach to identify candidate human CHD genes. Using human exome sequence data, mouse transcriptome data, and protein-protein interaction network analyses, we identify a subnetwork of potential CHD genes that includes multiple proteasome factor genes. We use CRISPR screening in zebrafish to identify roles for these proteosome factor genes in zebrafish heart development. Finally, we generate stable zebrafish mutant lines for two of the identified proteasome genes and use them to demonstrate novel functions for proteosome genes in heart development.

## Results

### Identification of 245 candidate CHD genes

To take a new approach to identifying candidate CHD genes, we turned to the high *s*_het_ genes, which are associated with Mendelian disorders as well as cellular and embryonic viability [23]. With specific respect to heart development, we find that genes with damaging *de novo* variants in patients with CHDs are strongly correlated with high *s*_het_ values, whereas no correlation is observed between *s*_het_ values and genes with damaging *de novo* variants in control subjects (Fig. 1A). In addition, most known CHD genes, as defined by Yang et al. (S1 Table; [22]), are found in the top *s*_het_ deciles (Fig. 1B), further implicating high *s*_het_ genes in CHDs.

**Fig. 1.**
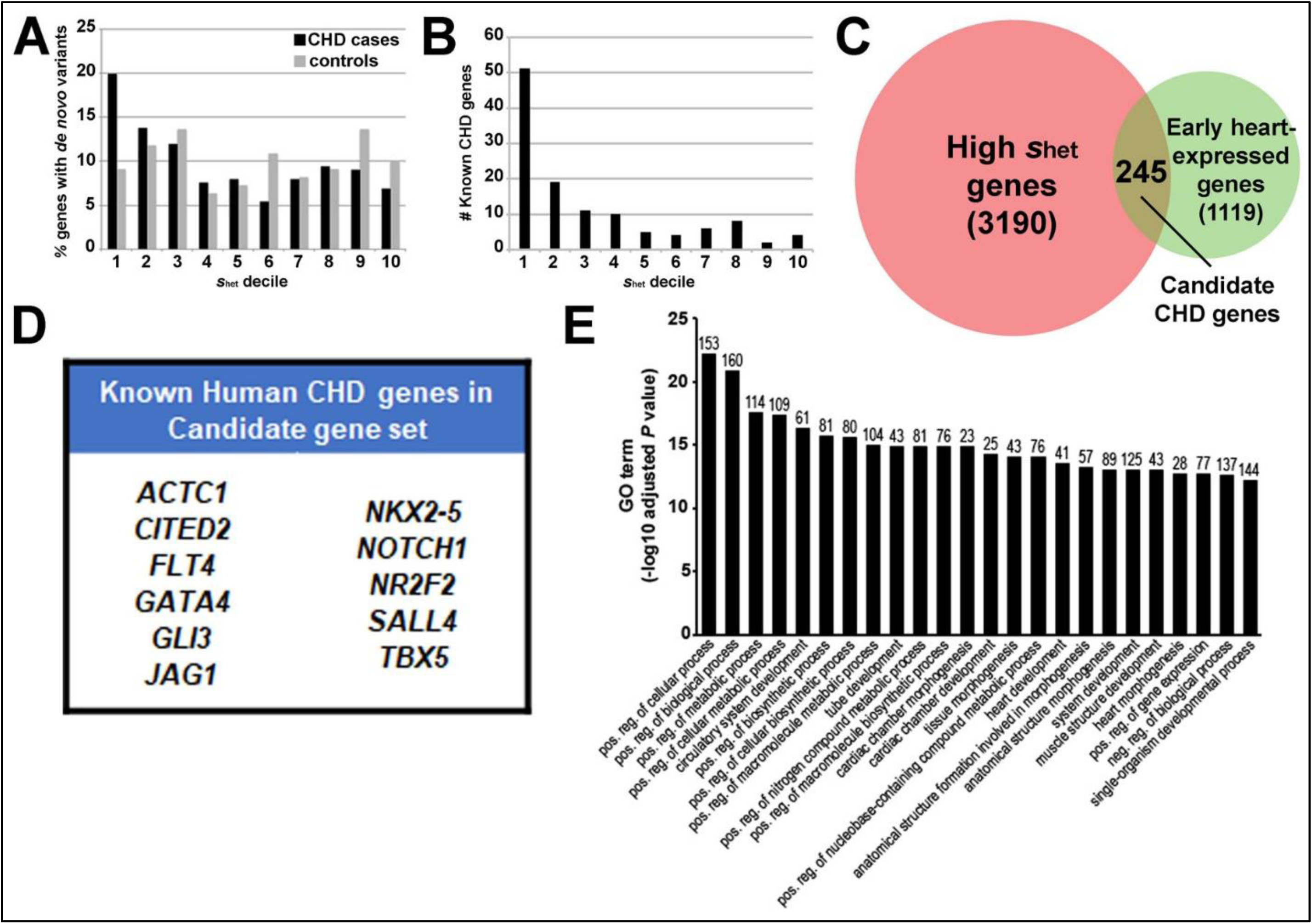
Identifying new candidate CHD genes. **A.** Correlation of high *s*_het_ genes with genes exhibiting damaging *de novo* variants in CHD cases (black bars; P=1.09E-19). No correlation is observed between *s*_het_ deciles and damaging *de novo* variants observed in controls (grey bars; P=0.7). *s*_het_ deciles are from Cassa et al.’s Supplemental Table 1 [23]. *de novo* variant data are from Jin et al.’s Supplemental Table 9 (CHD cases) and Supplemental Table 10 (controls) [18]. **B.** Correlation of high *s*_het_ genes with known CHD genes, defined by [22] (S1 Table). *s*_het_ deciles are from [23]. **C.** Venn diagram showing overlap of genes with high *s*_het_ values in pink (3190 genes; S2 Table; top 2 deciles in dataset from [23]) and genes expressed in the mouse cardiac muscle cell lineage in green (1119 genes; S3 Table; datasets from [33] and [34]). The intersection is 245 candidate genes for CHDs (S4 Table). **D.** 11 known human CHD genes in our set of 245 candidate genes. **E.** Gene Ontology (GO) term enrichment for candidate gene set, obtained using DAVID. The number of genes (out of 245) represented by each term is shown on top of each bar. Bonferroni-adjusted P values are shown.

We selected the genes with the highest *s*_het_ values (in the top two deciles, 3190 genes; S2 Table; [23]). Over half of known CHD genes are represented within these highest *s*_het_ gene cohorts (Fig. 1B). We asked which of the 3190 high *s*_het_ genes are likely to play a role in heart development by identifying those that are also expressed in the cardiac muscle cell lineage during embryonic development. For this analysis, we used cardiac gene expression datasets from two studies: the cardiac muscle cell lineage, obtained from single-cell RNA-seq of whole mouse embryos (cell cluster 34, stages E9.5-E13.5; [33]), and the differentially-expressed gene sets from single-cell RNA-seq of *Nkx2-5*-positive and *Isl1*-positive cells from E7.5-E9.5 mouse embryos [34]. We chose these datasets because they encompass early stages of mammalian heart development, they are comprised of early myocardial cell types, and genes with high levels of embryonic heart expression are enriched for mutations in CHD patients in exome studies [14,15,18,19]. These combined mouse gene expression datasets provide a cardiac muscle cell lineage gene set (1119 genes; S3 Table). The intersection of the early cardiac muscle-expression genes with the high *s*_het_ genes identified 245 genes that we termed our candidate CHD genes (Fig. 1C; S4 Table).

Of our 245 candidate genes, 11 are known CHD genes in humans (Fig. 1D). Among these 11 genes are well-characterized genes with respect to mammalian heart development and CHDs, including *GATA4*, *NKX2-5*, and *TBX5* [35,36]. We also find that 44 of our 245 genes have had potential damaging mutations identified in CHD cases [18] (S5 Table). We next employed a Gene Ontology (GO)-term analysis of our candidate gene list. While many of the top Biological Process GO terms enriched in our candidate gene list include general terms such as positive regulation of cellular and metabolic processes, several cardiac development and morphogenesis terms are also enriched in our candidate gene set (Fig. 1E). Furthermore, using published transcriptome data [37], we determined that 154 of our 245 genes are expressed in human embryonic heart tissues (S6 Table). Thus, the intersection of high *s*_het_ genes and cardiac cell-lineage gene expression identifies known as well as potential key players in heart development and CHDs, without reference to their known phenotype.

### Protein-Protein Interaction networks reveal potential new CHD gene modules

To further explore the relationships between our 245 candidate CHD genes and known CHD genes, we turned to the STRING online database [38] to generate a Protein-Protein Interaction (PPI) network. We created a large PPI consisting of 366 proteins that correspond to the 245 candidate genes and 132 known CHD genes, with 11 overlapping genes (Fig. 2A). This large network shows that proteins encoded by the candidate CHD genes share many functional and/or physical associations with known CHD proteins.

**Fig. 2.**
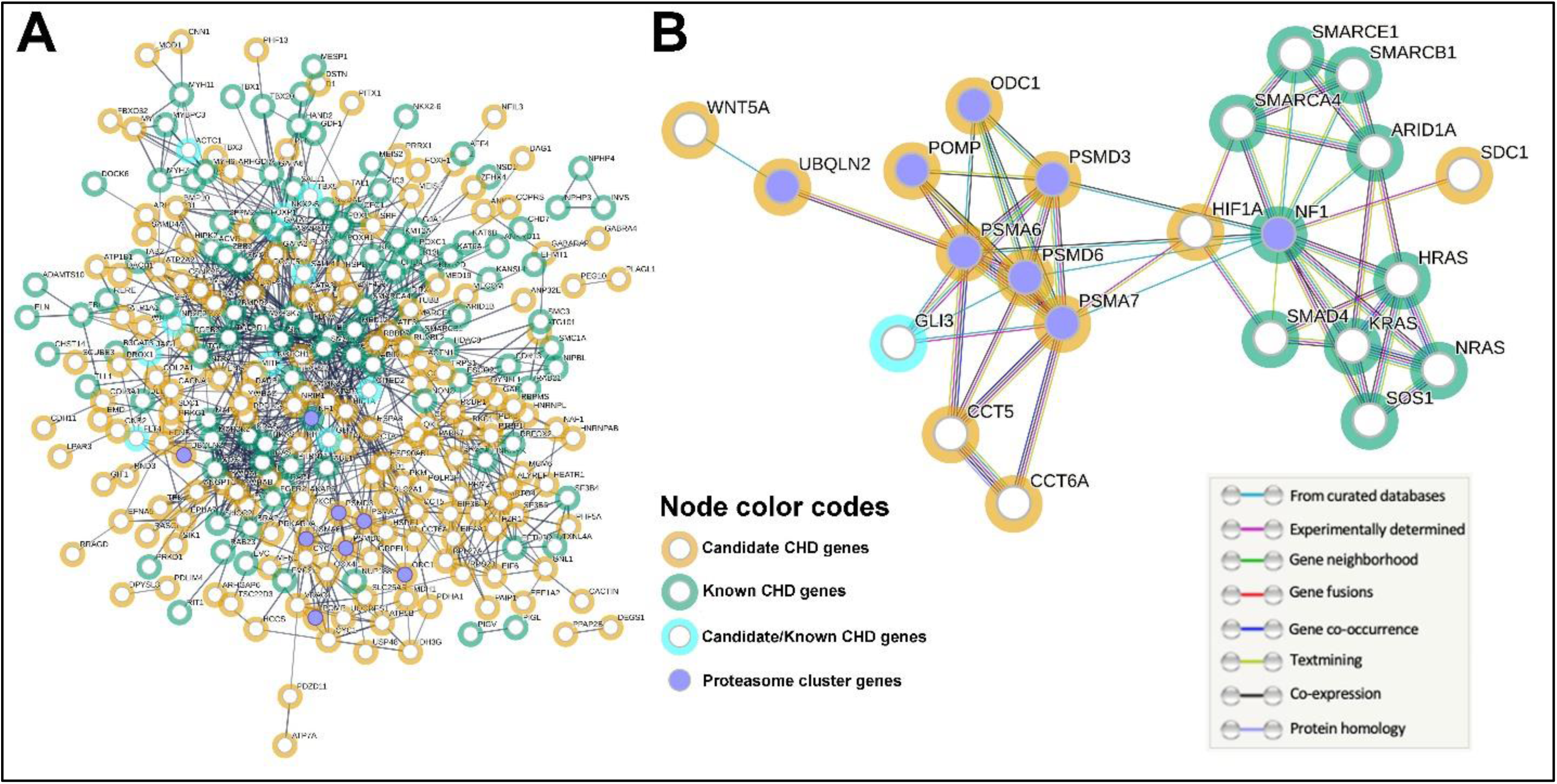
PPI networks reveal connections between known and candidate CHD genes. **A.** A Protein-Protein Interaction (PPI) network showing the evidence-based interactions among the proteins encoded by our 245 candidate CHD genes (yellow halos), 132 known CHD genes (green halos), and candidate genes that are also known CHD genes (blue halos, 11 genes from Fig. 1D). Interactions between proteins/nodes are indicated by a single grey line, whose thickness corresponds to the strength of the supporting data. This network consists of 365 proteins/nodes, contains 980 edges, and is associated with a PPI enrichment p-value < 1.0E-16. Purple nodes (Proteasome cluster genes) indicate the 8 factors (NF1, ODC1, POMP, PSMA6, PSMA7, PSMD3, PSMD6, and UBQLN2;) identified in the “Regulation of ornithine decarboxylase, and proteasome assembly” local cluster analysis (S7 Table). **B.** PPI subnetwork showing the interactions among the Proteasome cluster genes (purple nodes) and their direct connections from the larger PPI network in A. The node color code scheme is the same as in A. The lines between nodes are color coded to indicate the type of evidence supporting the protein-protein interaction. This subnetwork consists of 23 nodes, contains 58 edges, and has a PPI enrichment p-value < 1.0E-16.

To further assess the interactions embedded within our large network, we utilized STRING’s local network cluster analysis feature for functional enrichment. STRING identified 26 clusters, or functionally-enriched subnetworks (S7 Table). Most of these clusters consist mainly of proteins encoding the known CHD genes (S7 Table). The cluster with the highest percentage and number of candidate, but unproven, CHD genes is the “Regulation of ornithine decarboxylase, and proteasome assembly” cluster, or Proteasome cluster (S7 Table). The eight network genes in this cluster are *NF1*, *ODC1*, *POMP*, *PSMA6*, *PSMA7*, *PSMD3*, *PSMD6*, and *UBQLN2*, and these are highlighted in purple in the PPI network (Fig. 2A; S7 Table). *NF1* is a known CHD gene, while the remaining seven genes are candidate CHD genes. *POMP*, *PSMA6*, *PSMA7*, *PSMD3*, *PSMD6*, and *UBQLN2* all encode subunits of the proteasome complex or proteasome-interacting factors [39], and these factors have not previously been shown to cause CHDs.

To more closely examine the relationships of these eight Proteasome cluster-related factors, we used STRING to generate a subnetwork of these factors along with their direct connections within our larger PPI network. This subnetwork consists of 23 proteins encoded by 10 known, 12 candidate, and 1 overlapping candidate/known genes (Fig. 2B). This subnetwork illustrates the close connections of the proteasome factors PSMA7, PSMA6, PSMD6, and PSMD3 with known CHD factors, in particular NF1 and GLI3 (Fig 2B). Together, these findings support potential roles for our candidate CHD genes, in particular the Proteasome cluster genes, in heart development and CHDs.

### CRISPR knock downs in zebrafish embryos reveal roles for Proteasome cluster genes in heart development

We next wanted to test for roles for the Proteasome cluster genes in heart development by using CRISPR knock downs in zebrafish embryos. Six of these eight human genes, *NF1*, *ODC1*, *POMP*, *PSMA6*, *PSMD3*, and *PSMD6,* have orthologs in the zebrafish genome, with *NF1* and *PSMA6* having duplicate zebrafish orthologs: *nf1a* and *nf1b*, and *psma6a* and *psma6b* [40]. To test the functions of zebrafish *nf1a*, *nf1b*, *odc1*, *pomp*, *psma6a*, *psma6b*, *psmd3*, and *psmd6*, we turned to an efficient CRISPR knock down approach, using a pool of four CRISPR guide RNAs per gene [31]. As a positive control for using this approach to detect heart defects, we knocked down the zebrafish *hand2* gene with a pool of four *hand2* guide RNAs. *HAND2* is a known CHD gene [41,42] (S1 Table), and zebrafish *hand2* mutants show severe defects in early myocardial precursor cell migration, leading to two separate myocardial domains, or cardia bifida [43]. Zebrafish *hand2* CRISPR knock-down (CRISPR-KD) embryos show severe cardia bifida, observed using expression of the pan-myocardial transgene *myl7*:EGFP (Fig. 3A-B). Almost 100% of *hand2* CRISPR-KD embryos show severe bifida (Fig. 3C), thus closely resembling null *hand2^−/−^* mutant embryos [43]. These findings support the efficacy of the four-guide approach in attaining gene knock down and high frequency CRISPR-KD heart defects.

**Fig. 3.**
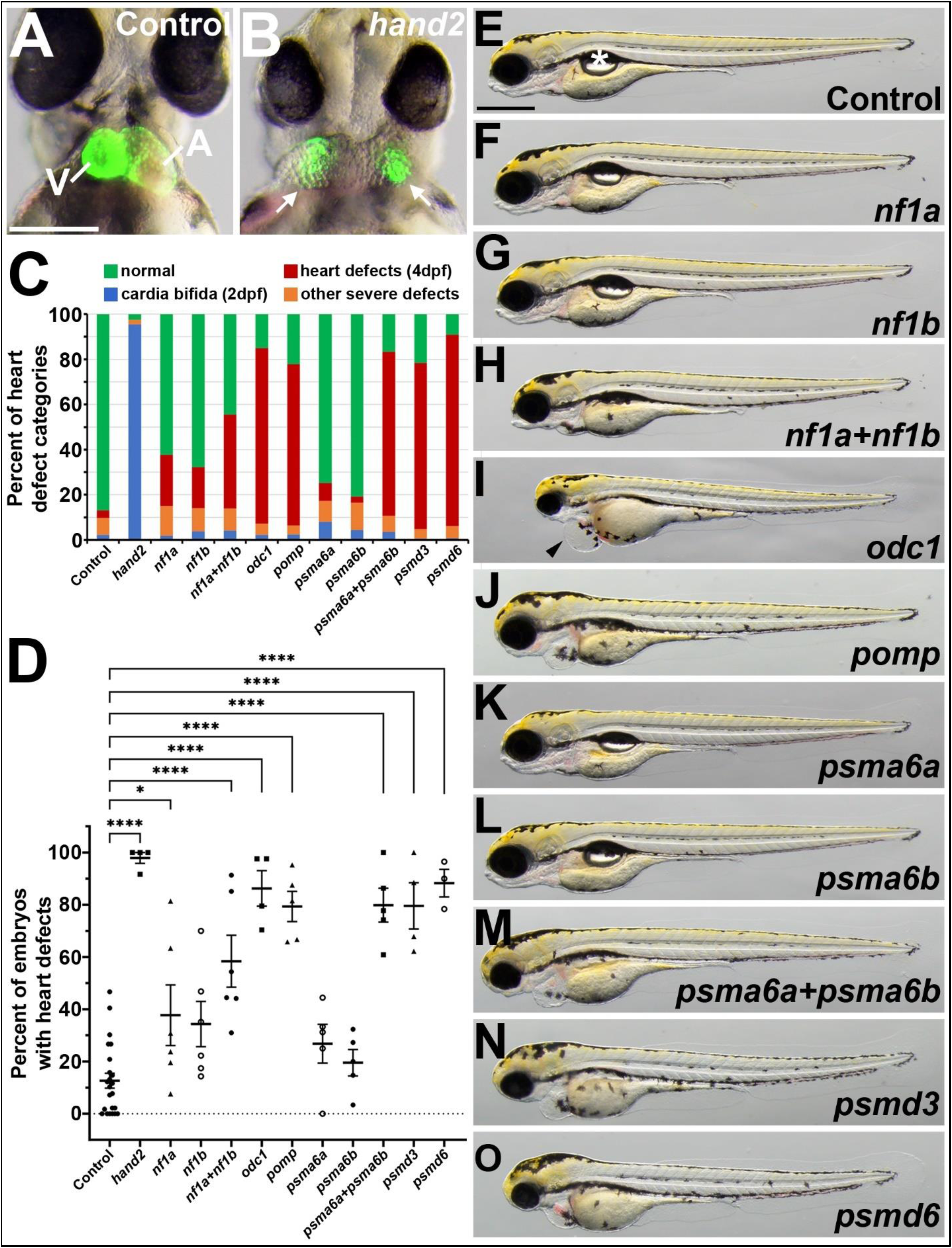
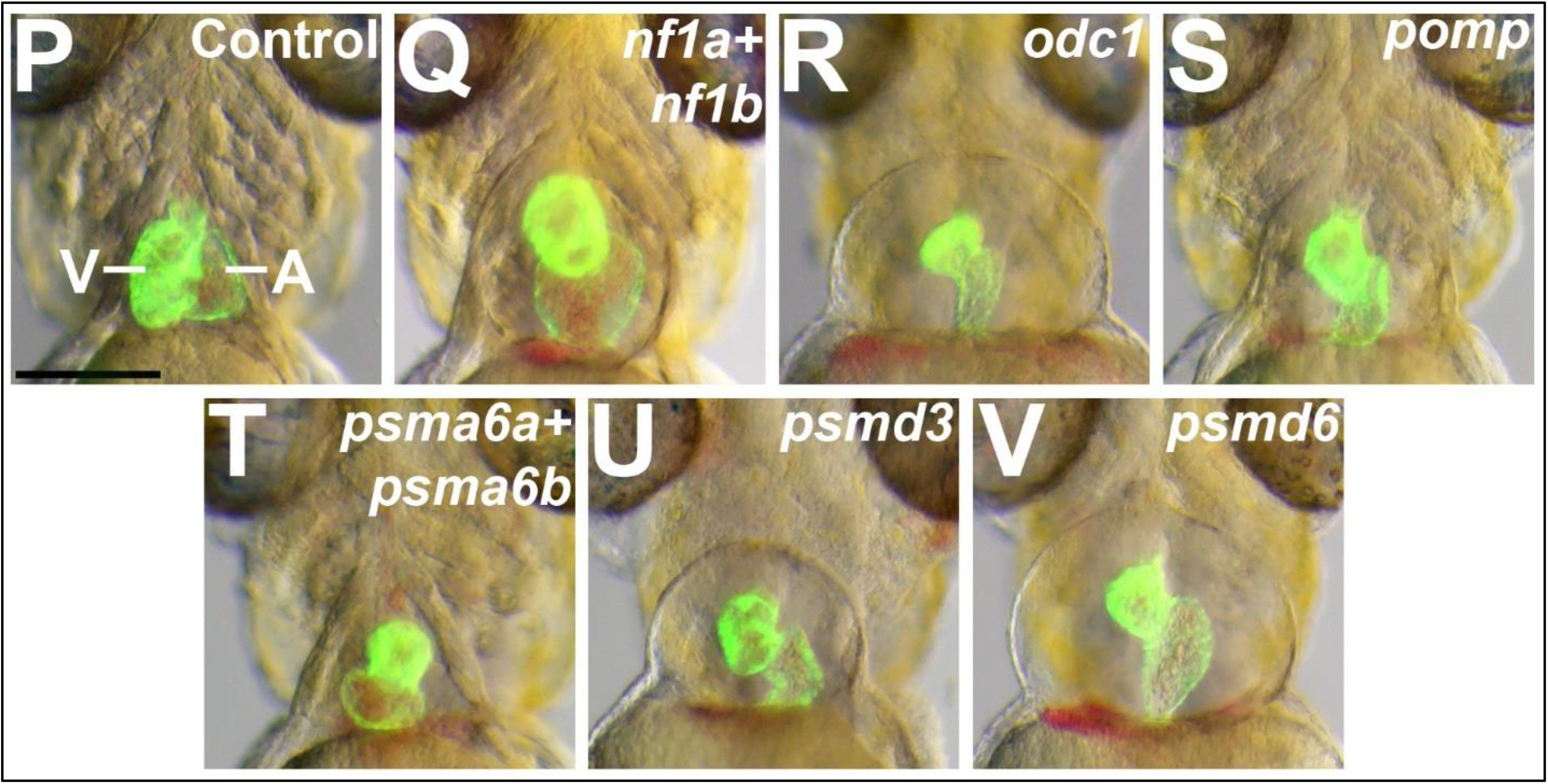
CRISPR knock downs (CRISPR-KD) of Proteasome cluster genes lead to heart defects in zebrafish embryos and larvae. **A.-C**. *hand2* CRISPR-KD leads to cardia bifida. **A.** 2 dpf control embryo showing *myl7:*EGFP heart. V, ventricle. A, atrium. Scale bar=200 μm. **B.** 2 dpf *hand2* CRISPR-KD embryo showing two domains (cardia bifida) of *myl7:*EGFP (arrows). **C.** Graph showing frequencies of different heart defect categories observed in CRISPR-KD embryos. Cardia bifida and other heart defects, such as heart looping defects (“other severe defects” in the legend), were scored at 2 dpf. Hearts were also scored at 4 dpf for defects arising between 2 dpf and 4 dpf (“heart defects (4dpf)” in the legend). Examples of these later-arising defects are illustrated in panels P-V. **D.** Graph showing frequencies of embryos with heart defects (at 4 dpf; all categories combined). Each dot represents an experimental replicate batch of embryos. N=3-7 replicates per gene of interest; N=24 for the control. Each replicate batch consists of 7-75 embryos (mean=38±16). * P<0.01. **** P<0.0001. **E.-O.** Lateral views of 4 dpf larvae, showing representative CRISPR-KD phenotypes. The bubble (asterisk in E) is the swim bladder, an indicator of healthy larvae [40]. The arrowhead in I points to severe heart cavity swelling/edema. Scale bar=500 μm. **P.-V.** Ventral views of hearts labeled with *myl7:*EGFP in 4 dpf larvae, showing representative CRISPR-KD heart phenotypes for genes shown. Scale bar=200 μm. V, ventricle. A, atrium.

We then used this CRISPR-KD approach to knock down the Proteasome cluster genes. Only a low percentage of embryos were observed to have cardia bifida or other heart defects at 2 days post fertilization (dpf) after knocking down the Proteasome cluster genes, similar to the incidence of these defects observed with control CRISPR (Fig. 3C). Knock down of *nf1a*, *nf1b*, *psma6a*, or *psma6b* individual genes did not lead to a high frequency of heart defects (Fig. 3C-D). However, at four days post-fertilization (4 dpf), knock-down of *nf1a+nf1b*, *odc1*, *pomp*, *psma6a+psma6b*, *psmd3*, and *psmd6* all led to a high frequency of defects in heart development, arising between 2 dpf and 4 dpf (Fig. 3C-D). In particular, these larvae show edema, or swelling around the heart cavity, and hearts in which the chambers appear malformed and not properly looped (Fig. 3E-V). The high frequency of heart defects in these CRISPR-KDs suggest that these Proteasome cluster genes play roles in heart development.

### *pomp* and *psmd6* mutant zebrafish embryos exhibit heart defects and other phenotypes

To further examine the roles for these proteasome factors in heart development, we generated stable mutant strains for *pomp* and *psmd6*. For both genes, we used CRISPR to generate alleles that delete the transcription start site (TSS) and 5’end of each gene (Fig. 4A-B). These TSS deletion alleles are *pomp^scm41^* and *psmd6^scm40^*, hereafter referred to as *pomp* and *psmd6* mutants.

**Fig. 4.**
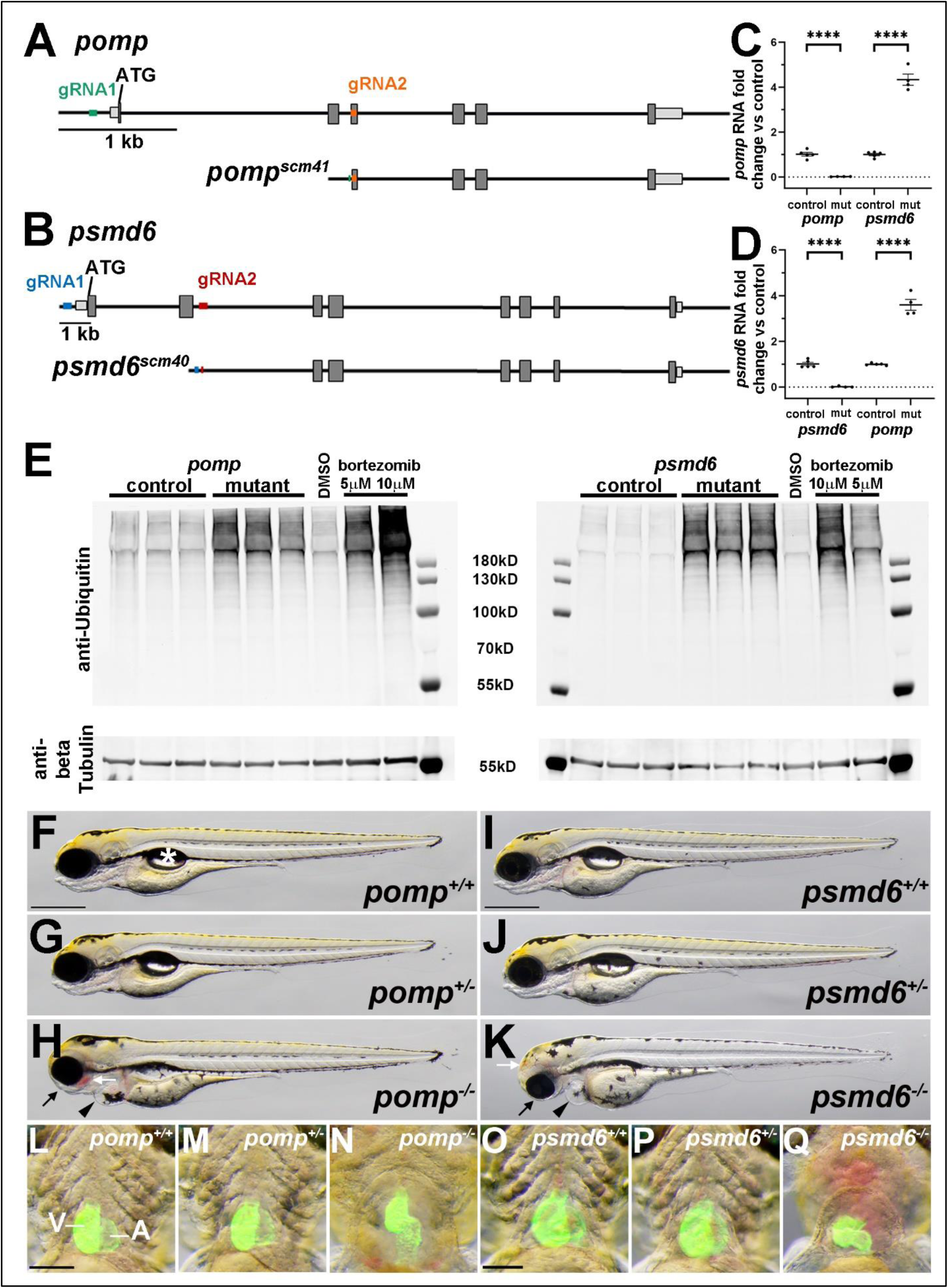
*pomp* and *psmd6* mutant zebrafish larvae show cardiac and extracardiac phenotypes. **A.** Schematic illustrating *pomp* gene structure and CRISPR deletion. *pomp* exons are shown as boxes and the start codon is labeled. Guide RNA target sites are labeled in green and orange. DNA sequencing of the *pomp^scm41^* strain showed the *pomp* gene sequence between the two guide RNA sites has been deleted. **B.** Schematic illustrating *psmd6* gene structure and CRISPR deletion. *psmd6* exons are shown as boxes and the start codon is labeled. Guide RNA target sites are labeled in blue and red. DNA sequencing of the *psmd6^scm40^* strain showed the *psmd6* gene sequence between the two guide RNA sites has been deleted. **C.-D.** qRT-PCR analysis of (C) *pomp* and (D) *psmd6* expression in 4 dpf phenotyped *pomp* and *psmd6* control (+/+ and +/-) or mutant (mut, −/−) larvae. Each dot represents a replicate batch of embryos. N=5 (control) or 4 (mutant) replicates. Each replicate consists of 10-20 embryos. **** P<0.0001. **E.** Western analysis using anti-Ubiquitin and anti-beta Tubulin (as a loading control). Larvae from *pomp* and *psmd6* clutches were phenotyped as control (+/+ and +/-) or mutant (−/−) larvae at 4 dpf and collected for lysates. Lysates were prepared from 3 replicate pools of animals for each condition, with n = 20 animals per replicate. Larvae from bortezomib or DMSO control treatments were collected at 4 dpf, with n=20 animals per lysate. Molecular weight markers are shown. **F.-K.** Images of live 4 dpf larvae. The asterisk in F marks the swim bladder. Arrowheads in H, K point to heart cavity swelling/edema. Black arrows in H, K point to reduced craniofacial structures. White arrows in H, K point to areas of blood pooling. N= 3 *pomp^+/+^*, 12 *pomp^+/-^*, 9 *pomp^−/−^*, 4 *psmd6^+/+^*, 8 *psmd6^+/-^*, 6 *psmd6^−/−^*. Scale bars=500μm. **L.-Q.** Ventral views of hearts labeled with *myl7:*EGFP in 4 dpf larvae. N= 7 *pomp^+/+^*, 9 *pomp^+/-^*, 7 *pomp^−/−^*, 3 *psmd6^+/+^*, 9 *psmd6^+/-^*, 5 *psmd6^−/−^*. Scale bars=200 μm. V, ventricle. A, atrium.

To confirm that these deletion alleles cause loss of expression of their respective genes, we used qRT-PCR to examine *pomp* and *psmd6* expression. As expected, *pomp* expression is lost in *pomp* mutant larvae, and *psmd6* expression is lost in *psmd6* mutant larvae (Fig. 4C-D). We observe that *pomp* expression is upregulated in *psmd6* mutant larvae, and *psmd6* is upregulated in *pomp* mutant larvae (Fig. 4C-D), indicating a compensation effect on gene expression for these proteasome factors. To confirm that loss of *pomp* and *psmd6* functions lead to a defect in proteasome function, we examined levels of ubiquitinated proteins. We observe an increase in ubiquitinated proteins in both *pomp* and *psmd6* mutant larvae (Fig. 4E). This increase is similar to that observed when wild-type embryos are treated with the proteasome inhibitor bortezomib (Fig. 4E). These results show that we have generated transcription-null alleles of *pomp* and *psmd6* and that loss of these genes leads to the expected defects in proteasome function.

To examine the phenotypes of *pomp* and *psmd6* mutant embryos, we used incrosses of heterozygous *pomp* and *psmd6* fish. *pomp^+/+^*, *pomp^+/-^*, *psmd6^+/+^,* and *psmd6^+/-^*larvae exhibit normal body and heart morphology at 4 dpf (Fig. 4F,G,I,J,L,M,O,P). For both *pomp* and *psmd6* homozygous mutants, we observed heart edema and heart morphology defects by 4 dpf, similar to what we observed with *pomp* and *psmd6* CRISPR knockdowns (Fig. 4H,K,N,Q). We also observed additional phenotypes in these mutant larvae, including reduced craniofacial structures and blood pooling in the head (Fig. 4H,K). These results support our findings from the CRISPR knockdown tests and provide support for the Proteasome cluster genes playing similar roles in heart development.

### *pomp* and *psmd6* mutant hearts exhibit cellular blebbing

We next investigated when heart development issues arise in *pomp* and *psmd6* mutant embryos. We collected embryos at a series of time points during embryonic development and examined expression of the pan-cardiomyocyte and myocardial marker *myl7*, using RNA *in situ* hybridization and *myl7:*EGFP [44,45]. At 18 hpf, *myl7* is expressed in cardiomyocyte precursors in the anterior lateral plate mesoderm (ALPM) in control embryos, and this expression appears normal in *pomp* and *psmd6* mutants (Fig. 5A-C). At the time of myocardial tube formation at 24 hpf, *myl7* expression and tube formation appear normal in *pomp* and *psmd6* mutants (Fig. 5D-F). At the stage of early chamber formation and heart looping (48 hpf), and continuing to about 72 hpf, *myl7* expression and heart morphology continue to appear largely normal in *pomp* and *psmd6* mutants (Fig. 5G-L). These findings suggest that initial heart development occurs normally in *pomp* and *psmd6* mutants.

**Fig. 5.**
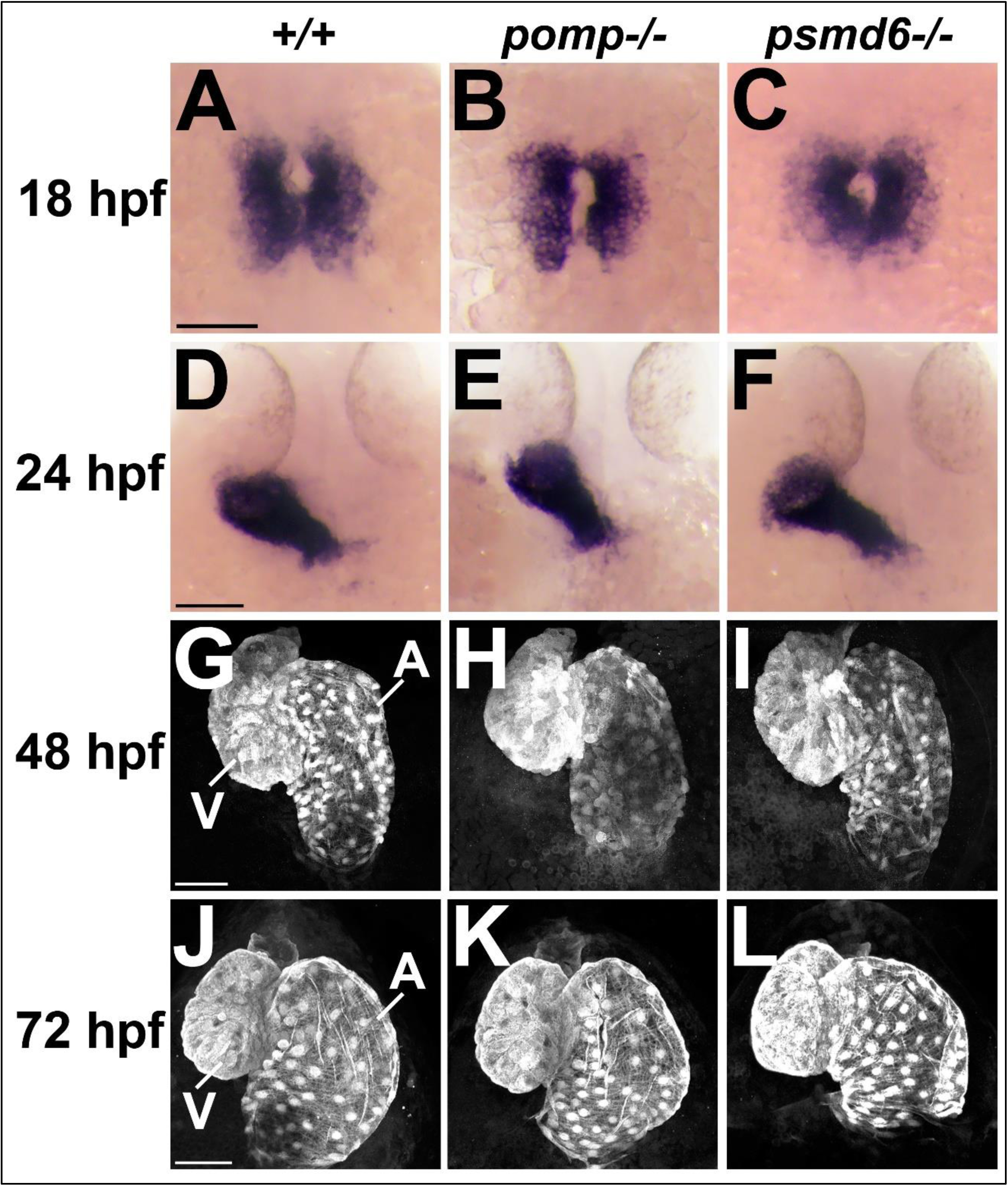
Early heart development appears normal in *pomp* and *psmd6* mutants. **A.-C.** Dorsal views of ALPM labeled for *myl7* in blue in 18 hpf embryos. N= 6 *pomp^+/+^*, 7 *psmd6^+/+^*, 11 *pomp^−/−^*, 8 *psmd6^−/−^*. Scale bar=100 μm. **D.-F.** Dorsal views of hearts labeled for *myl7* in blue in 24 hpf larvae. N= 8 *pomp^+/+^*, 8 *psmd6^+/+^*, 6 *pomp^−/−^*, 10 *psmd6^−/−^*. Scale bar=100 μm. **G.-I.** Ventral views of hearts labeled with *myl7:*EGFP in 48 hpf embryos. N= 16 *pomp^+/+^*, 12 *psmd6^+/+^*, 14 *pomp^−/−^*, 12 *psmd6^−/−^*. Scale bar=50 μm. V, ventricle. A, atrium. **J.-L.** Ventral views of hearts labeled with *myl7:*EGFP in 72 hpf larvae. N= 38 *pomp^+/+^*, 24 *psmd6^+/+^*, 45 *pomp^−/−^*, 29 *psmd6^−/−^*. Scale bar=50 μm. V, ventricle. A, atrium.

We then examined 4 dpf hearts using *myl7:*EGFP. Maximum intensity projections of the entire heart made at lower magnification have the effect of highlighting the ventral surface morphology due to its greater brightness relative to the deeper tissue. By examining these images, we observed abnormal arrangement of the atrium and ventricle and rounding-up of the cells on the myocardial surface in both *pomp* and *psmd6* mutants (Fig. 6A-C). By examining projections of a few slices obtained at higher magnification through the ventral ventricular wall, we observed that the trabecular myocardial cells also appeared rounded (Fig. 6D-F). To quantify this cellular blebbing phenotype, we counted the number of rounded cells per heart.

**Fig. 6.**
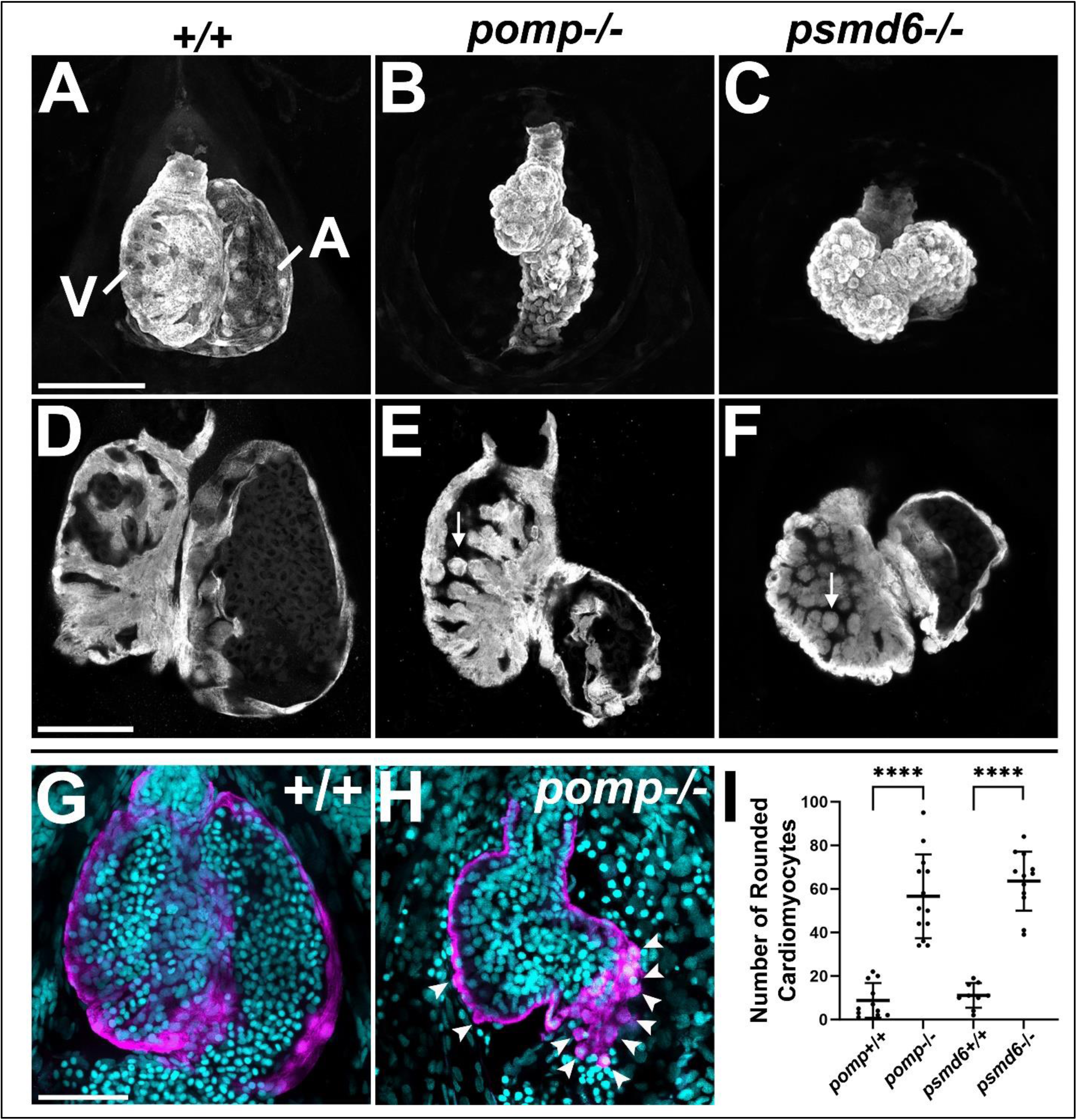
*pomp* and *psmd6* mutants exhibit myocardial cell blebbing. **A.-C.** Ventral views of hearts as maximum intensity projections (MIPs) of the entire myocardium, labeled with *myl7:*EGFP in 4 dpf larvae. Presence of dysmorphic cardiac chambers: 1/17 *pomp^+/+^*, 0/15 *psmd6^+/+^*, 30/30 *pomp^−/−^*, 27/27 *psmd6^−/−^* larvae. Morphology of mutant hearts: *pomp^−/−^* larvae show 24/30 elongated hearts (as in B) and 6/30 lateral hearts; *psmd6^−/−^* larvae show 2/27 elongated hearts and 25/27 lateral hearts (as in C). Scale bar=100 μm. V, ventricle. A, atrium. **D.-F.** Ventral views of hearts as MIPs of 2 to 7 slices (1.6 – 5.6μm total section thickness) through the ventral ventricular wall, labeled with *myl7:*EGFP in 4 dpf larvae. Arrows in E and F indicate examples of rounded cardiomyocytes in the interior wall of the ventricle. Hearts in D – F are from different larvae than those in A – C. Presence of rounded trabecular cardiomyocytes: 0/17 *pomp^+/+^*, 1/15 *psmd6^+/+^*, 25/29 *pomp^−/−^*, 24/27 *psmd6^−/−^*. Scale bar=50 μm. **G.-H.** Ventral views of hearts labeled with *myl7:*EGFP (magenta) and DAPI (cyan) in 4 dpf larvae. Arrowheads in (H) point to rounded cardiomyocytes protruding outward from the myocardium. Scale bar=50 μm. **I.** Graph showing numbers of rounded cardiomyocytes per heart at 4 dpf. Only cells clearly projecting outward from the surface of the heart were counted. Each dot represents a heart/larva. N=9-13 per genotype. **** P<0.0001.

Unambiguously identifying individual rounded cells within the ventricle proved difficult, so we focused on counting cardiomyocyte nuclei (double-positive for GFP and DAPI) that clearly protruded outward from the heart wall. While wild-type hearts occasionally showed such rounded cells, *pomp* and *psmd6* mutant hearts showed about 5X more rounded cells (Fig. 6G-I).

### *pomp* and *psmd6* mutant hearts exhibit reduced outflow tracts

We then examined additional markers associated with myocardial differentiation and heart development. To assess myocardial sarcomere formation in *pomp* and *psmd6* mutants, we examined immunostaining οf α-actinin, which localizes to Z-discs in striated cardiac and skeletal muscle [46,47]. We observe that α-actinin is organized in its periodic pattern in wild-type and in *pomp* and *psmd6* mutant hearts at 4 dpf (Fig. 7A-E). No significant differences in sarcomere lengths or myofibril widths were observed between wild-type and mutant hearts (Fig. 7F-G), suggesting that *pomp* and *psmd6* are not required for sarcomere or myofibril formation.

**Fig. 7.**
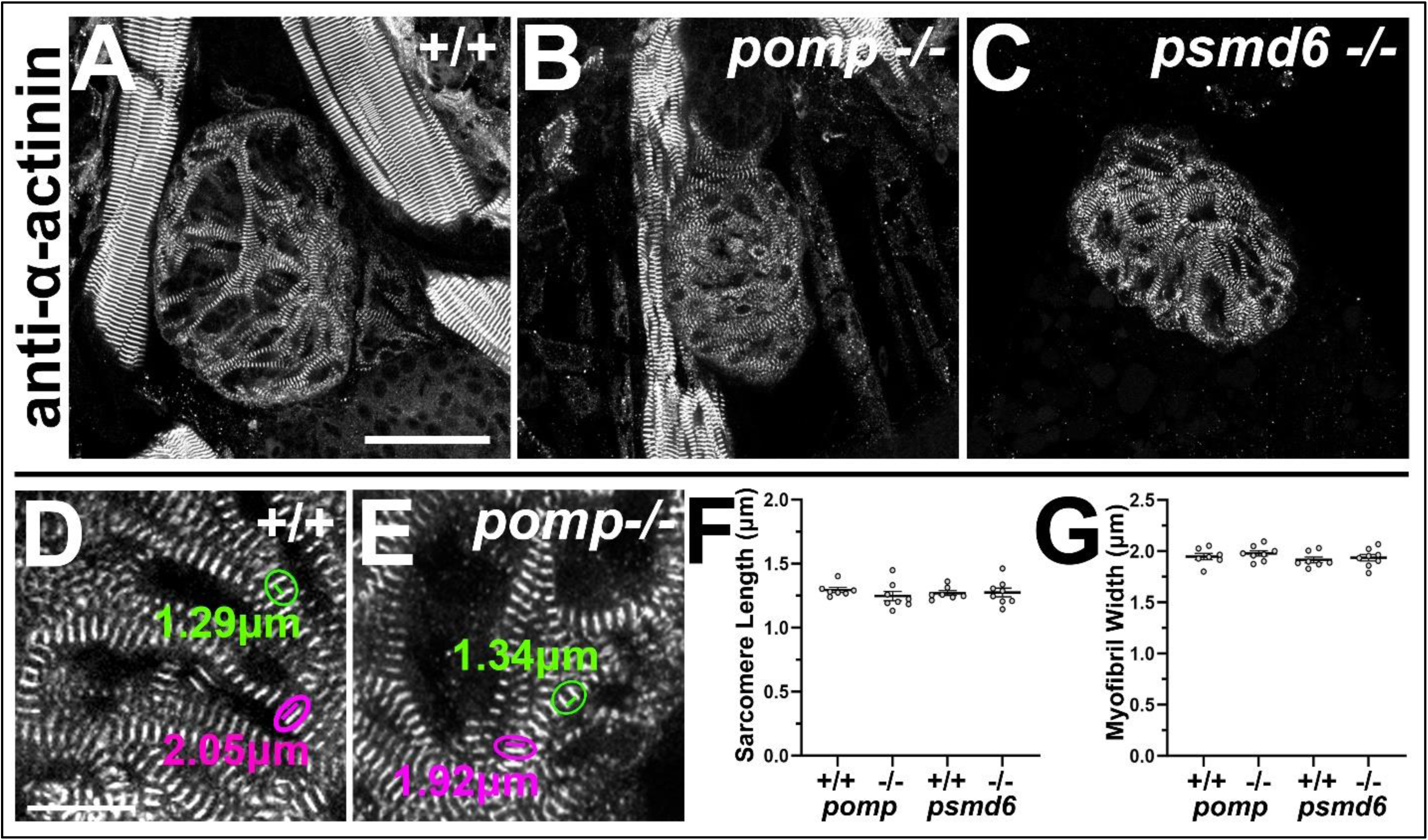
Myocardial sarcomere and myofibril formation appear normal in *pomp* and *psmd6* mutants. **A.-C.** Ventral views of ventricles labeled with α-actinin (A-C) in 4 dpf larvae. N= 5 *pomp^+/+^*, 7 *psmd6^+/+^*, 8 *pomp^−/−^*, 9 *psmd6^−/−^*. Scale bar=25 μm. **D.-E.** Magnified views of α-actinin stain. Encircled green lines illustrate sarcomere lengths, with corresponding measurements shown. Encircled magenta lines illustrate myofibril widths, with corresponding measurements shown. **F.-G.** Graphs showing (F) sarcomere length and (G) myofibril width measurements at 4 dpf. Each dot represents the average of 9-14 measurements (one per myofiber) from a single heart/larva. N=7 for *pomp^+/+^* and *psmd6^+/+^*. N=8 for *pomp^−/−^* and *psmd6^−/−^*. No significant differences were observed between wild-type and mutant hearts.

To assess formation of the outflow tract in *pomp* and *psmd6* mutants, we examined expression of the outflow tract smooth muscle marker *elastinb* (*elnb*) [48]. Early expression of *elnb* appears somewhat reduced in *pomp* and *psmd6* mutant hearts at 72 hpf (Fig. 8A-C). However, by 96 hpf, the *elnb* expression domain appears strongly reduced and dysmorphic (Fig. 8D-F). These results show that *pomp* and *psmd6* are needed for proper outflow tract formation.

**Fig. 8.**
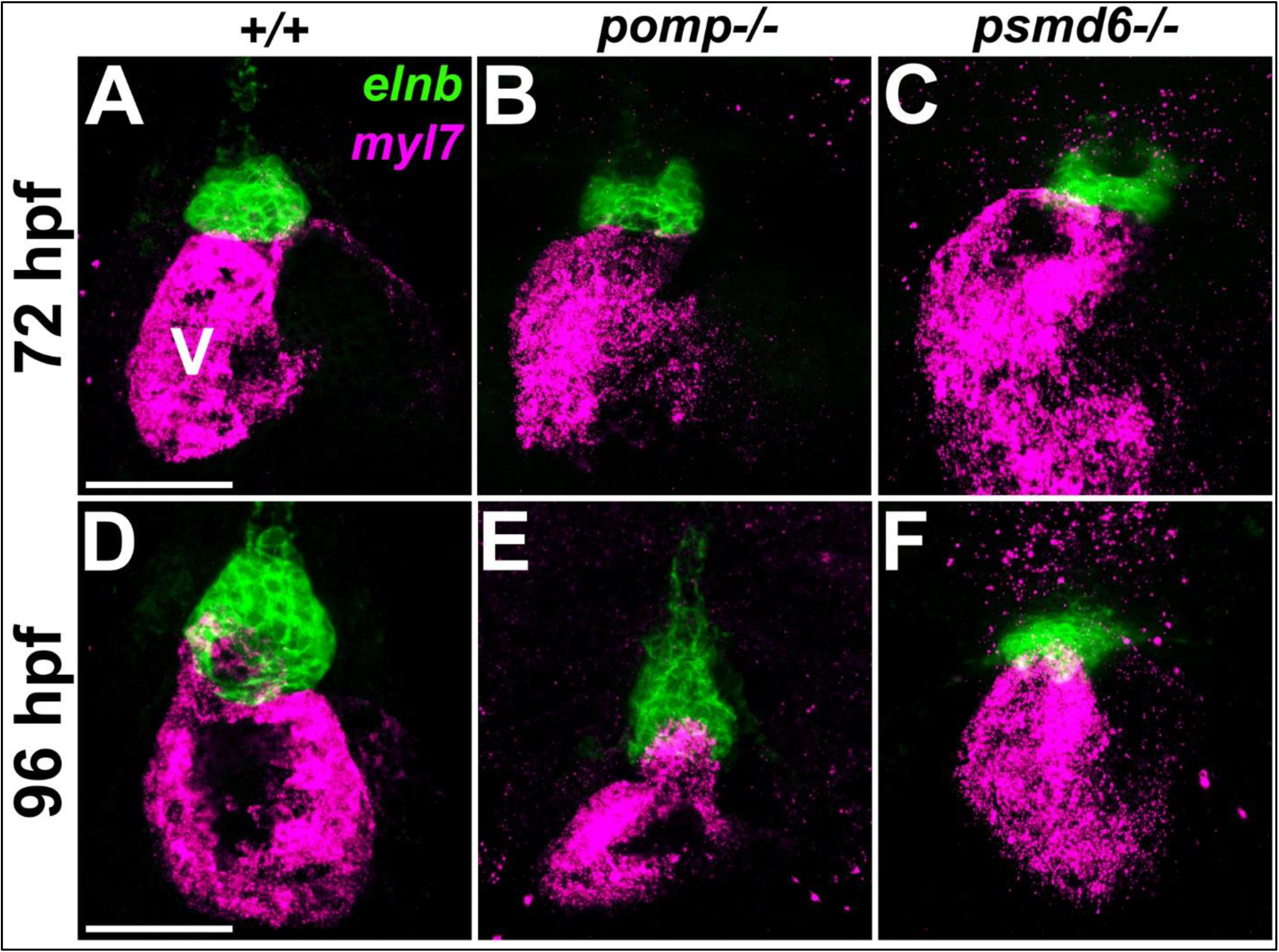
*pomp* and *psmd6* mutants exhibit reduced outflow tracts. **A.-C.** Ventral views of hearts co-labeled with *myl7* and *elnb* in 72 hpf larvae. Percent of embryos with reduced *elnb* expression (N): 22% (4/18) *pomp^+/+^*, 0% (0/15) *psmd6^+/+^*, 70% (23/33) *pomp^−/−^*, 72% (13/18) *psmd6^−/−^*. Scale bar=50 μm. V, ventricle. **D.-F.** Ventral views of hearts co-labeled with *myl7* and *elnb* in 96 hpf larvae. Percent of embryos with reduced *elnb* expression (N): 0% (0/6) *pomp^+/+^*, 0% (0/18) *psmd6^+/+^*, 47% (7/15) *pomp^−/−^*, 100% (7/7) *psmd6^−/−^*. Scale bar=50 μm.

## Discussion

In this study, we take a novel approach to identify candidate human CHD genes. Using multiple data systems, including human exome sequences, mouse transcriptomes, and protein-protein interactions, we identify a subnetwork of potential CHD genes that includes multiple genes encoding proteasome factors. We use CRISPR screening in zebrafish to demonstrate roles for these proteosome factor genes in zebrafish heart development. Furthermore, our analyses of stable mutant zebrafish strains for two of these genes, *pomp* and *psmd6*, reveal novel roles for proteosome genes in heart development. Our work addresses a major hurdle for understanding the causes of CHD, through the identification and validation of a new set of candidate CHD genes.

A major innovative aspect of this study is our identification of a set of 245 human genes with presumptive roles in heart development and CHDs. The standard approach to identify CHD genes is through using exome and genome sequencing of CHD patients and their families to identify deleterious mutations. In contrast, our approach identifies candidate CHD genes through their lack of deleterious mutations in the normal human population. While our list of 245 candidate CHD genes includes several known CHD genes, most of the 245 genes have unknown requirements in early heart development. There are about 130 known CHD genes, but it is estimated that there are over 400 genes that contribute to CHDs [18,22]. Thus, our list of candidate CHD genes could represent a majority of unknown CHD genes.

We use PPI network analyses to identify a functionally-related subnetwork that includes five genes encoding proteasome factors: *POMP*, which encodes Proteasome maturation protein, and *PSMA6*, *PSMA7*, *PSMD3*, and *PSMD6*, which encode proteasome subunits. The proteasome is a multi-subunit complex that is part of the ubiquitin-proteasome system (UPS), which is responsible for carrying out the majority of normal cellular protein degradation [39,49]. *PSMA6* and *PSMA7* both encode α subunits of the 20S core particle of the proteasome. *PSMD3* and *PSMD6* both encode non-ATPase subunits of the proteasome’s 19S regulatory particle. *POMP* (also known as Ump1) functions as an assembling chaperone for the 20S proteasome [39]. The proteasome and the UPS have well-established roles in adult cardiac diseases [50]. However, proteasome factors have thus far only been indirectly linked with CHDs. Only single, likely deleterious *de novo* variants in *POMP* and *PSMD6* have been identified in CHD exome studies [18] (see S5 Table). Mutations in *PSMD12*, which encodes a 19S non-ATPase subunit, have been linked with a neurodevelopmental syndrome that in some cases involves CHDs [51]. Roles for specific proteasome factors in heart development have not previously been addressed.

To demonstrate the functions of proteasome genes in early heart development, we use zebrafish CRISPR knock-downs and mutant strains. Zebrafish have many advantages for discovering and characterizing new CHD genes [28–32]. In our study, we take advantage of a previously-established CRISPR screening approach [31] to show that our subnetwork genes are all required for zebrafish heart development. Although some of the genes we test have duplicate copies in the zebrafish genome, we were able to use co-CRISPR-KD of gene duplicates to demonstrate their functions in heart development. Furthermore, using stable mutant strains, we show that *pomp* and *psmd6* mutant embryos have phenotypes that closely resemble, and thus validate, the CRISPR-KDs.

A key finding from our study is that zebrafish *pomp* and *psmd6* mutant embryos share heart development phenotypes that are relevant to human CHDs. One characteristic phenotype we observe is myocardial cell blebbing. This phenotype has also been observed in other zebrafish mutants for critical heart development genes, including *snai1b*, *tcf21*, *wt1a*. *flii*, and *klf2a;klf2b* double mutants [52–55]. In these examples, myocardial cell blebbing has been shown to be due to cell adhesion and cell polarity defects but not associated with increased cell death [52–55]. Another phenotype we observe in *pomp* and *psmd6* mutant embryos is defective outflow tract formation, which is also observed in many zebrafish mutants for heart development genes [28,40,56,57]. The zebrafish outflow tract shares evolutionary conservation with the mammalian outflow tract [58–60], and the outflow tract is a major site of human CHDs [61,62]. Further studies are needed to understand how proteasome factors might interact with known heart development genes in promoting myocardial cell adhesion and outflow tract formation.

A limitation of our study is that it is unclear how directly the proteasome factor genes are functioning in heart development. *pomp*, *psmd6*, and other genes encoding proteasome factors are very broadly expressed throughout zebrafish embryogenesis [40,63]. *pomp* and *psmd6* mutant embryos exhibit extracardiac defects, including reduced craniofacial development. These findings are consistent with the craniofacial defects observed upon loss of other zebrafish proteasome factors [51,63]. These issues make it challenging to identify the primary mechanism through which proteasome factors are needed for heart development. Future studies employing zebrafish genetic mosaics or mouse conditional mutant strains could help address when, and in which tissue(s), proteasome factors influence heart development.

## Materials and Methods

### Identifying high s_het_ genes expressed in early heart development

We took the top 2 deciles of *s*_het_ genes (3199 genes) from Cassa et al.’s Supplemental Table 1 [23]. We entered these 3199 gene symbols into the Multi-symbol checker in HGNC (HUGO Gene Nomenclature Committee site; https://www.genenames.org/tools/multi-symbol-checker/) [64]. We collected the updated approved gene symbols to generate a revised list of 3190 high *s*_het_ genes (S2 Table).

Mouse cardiac muscle cell lineage datasets were generated from [33,34]. Cao et al. [33] performed single-cell RNA sequencing on whole mouse embryos from E9.5-E13.5. We took all genes from Cao et al.’s Cluster 34 Cardiac Muscle Lineages (gene list obtained from the file DE_gene_main_cluster.csv, downloaded from https://oncoscape.v3.sttrcancer.org/atlas.gs.washington.edu.mouse.rna/downloads). We entered this list into the HGNC HCOP (Comparison of Orthology Predictions) tool to obtain 610 human genes. Jia et al. [34] performed single-cell RNA sequencing on isolated cardiac progenitor cells from E7.5-E9.5 mouse embryos. We combined the gene lists from Jia et al.’s Supplementary Data 1 (list of genes that are differentially expressed in 3 *Nkx2.5*-positive cardiac progenitor cell clusters) and Supplementary Data 2 (list of genes that are differentially expressed in 5 *Isl1*-positive cardiac progenitor cell clusters). We entered these combined lists into the HGNC HCOP tool to obtain 651 human genes. These two gene sets were then combined, with duplicates removed, to identify 1119 genes (S3 Table). We used BioVenn (https://www.biovenn.nl/) to identify the overlap between the high *s*_het_ genes (3190 genes) and genes expressed in the mouse cardiac muscle cell lineage (1119 genes) (Fig. 1C; S4 Table). Venn diagrams were generated using BioVenn.

BioVenn was used to determine the overlap between our set of 245 candidate CHD genes and 132 known CHD genes (Fig. 1D). BioVenn was also used to determine the overlap between the 245 Candidate CHD genes and genes exhibiting damaging *de novo* mutations in CHD cases (Supplemental Table 9 from [18]; S5 Table).

A list of early human cardiac-expressed genes was generated from [37], which performed spatial transcriptomics on hearts from 4.5-9 week-old human embryos and single-cell RNA sequencing of hearts from 6.5 week-old human embryos. We combined all differential expression gene lists from their Supplemental Table 2 and Supplemental Table 3 to generate a list of 4670 genes. BioVenn was used to determine the overlap between these 4670 genes and the 245 candidate CHD genes (S6 Table).

For functional annotation, the 245 candidate CHD gene list was loaded into the Functional Annotation tool in DAVID ([65]; DAVID Knowledgebase version 2023q2; originally DAVID v6.7 2020), selecting functional annotation category GOTERMS.

### Constructing Protein-Protein Interaction networks

To construct Protein-Protein Interaction (PPI) networks, we used STRING [38], initially using v11.0 but also using versions up to v12.0. To generate the large network, we created a color-coding effect utilizing the “Payload datasets” feature within STRING. To generate a new payload dataset, we selected Homo sapiens and entered our list of known, candidate, and overlapping (candidate/known) gene symbol names. We made three different color categorizations and assigned a color code to each gene/node within each category. We used the following settings for the large network: We selected high confidence 0.7 for the minimum required interaction score. We selected the “max number of interactions to show” to be “none” for both the 1^st^ and 2^nd^ shell, to ensure the only nodes in the network are those from our candidate and known CHD gene lists. We selected “confidence” for “meaning of network edges”. We selected all active interaction data sources. We selected “hide disconnected nodes in the network”.

We utilized STRING’s local network cluster analysis for functional enrichment. We selected cluster CL:2692 “Regulation of ornithine decarboxylase, and proteasome assembly” within the Analysis tool in order to highlight the nodes/genes associated with that cluster in our network.

To generate a subnetwork based on the network nodes in CL:2692, we generated a new gene list. We started with the 8 nodes identified in CL:2692, and we then used the node interaction data from our large network to identify all proteins within our large network that share a direct connection with any of the eight nodes. A subnetwork was generated with these genes as inputs, with nodes color-coded as for the large network. We again used a minimum required interaction score of 0.7, utilized all active interaction data sources, and limited the nodes included within our subnetwork to only those specifically inputted by setting the max number of interactions to show to none for both the 1^st^ and 2^nd^ shell. We selected “evidence” for “meaning of network edges”, with the line colors indicating the different types of evidence supporting the interactions between two given proteins.

### Zebrafish husbandry

All experiments involving live zebrafish (*Danio rerio*) were carried out in compliance with Seattle Children’s Research Institute’s Institutional Animal Care and Use Committee guidelines. Zebrafish were raised and staged as previously described [66,67]. Time indicated as hpf or dpf refers to hours or days post-fertilization at 28.5°C, respectively.

The wild-type stock and genetic background used was AB. The *Tg(myl7:EGFP)^twu34^* line has been previously described [45]. For fish stock maintenance, eggs were collected from 20–30 min spawning periods and raised in Petri dishes in ICS water [68] in a dark 28.5°C incubator, up to 5 dpf. After 5 dpf, the fish were maintained on a recirculating water system (Aquaneering) under a 14 h on, 10 h off light cycle. From 6–30 dpf, the fish were raised in 2.8 L tanks with a density of no more than 50 fish per tank and were fed a standard diet of paramecia (Carolina) one time per day and Zeigler AP100 dry larval diet two times per day. From 30 dpf onwards, the fish were raised in 6 L tanks with a density of no more than 50 fish per tank and were fed a standard diet of *Artemia nauplii* (Brine Shrimp Direct) and Zeigler adult zebrafish feed, each two times per day.

### CRISPR knock-downs in zebrafish embryos

The sequences for the oligonucleotides used to synthesize the single-guide RNAs for CRISPR-KDs were taken from the published genome-scale Lookup Table [31] and are provided in S8 Table. For a negative control 4-guide set, we used the “Genetic Screen Scramble1 Control” guides [31] (S8 Table). sgRNAs were synthesized as described [31]. Pools of four gene-specific (or control) oligos, each incorporating a T7 RNA polymerase site, were annealed to a common scaffold oligo and transcribed *in vitro* to generate pools of four sgRNAs, as described [31]. For CRISPR-KD phenotype analysis, 2 µL of a 4-guide cocktail of sgRNA at 2 µg/µL was combined with 2 µL of Cas9 protein (IDT Alt-R *S.p.* Cas9 Nuclease V3, Cat. # 1081058) at 10 µM (diluted as in [31]) and incubated at 37°C for 5 minutes. In cases where two 4-guide cocktails were combined, 1 µL of each cocktail was used. 1 µL of phenol red injection solution (0.1% phenol red and 0.2M KCl in water) was added to generate the working solution for embryo injections. 1-cell stage embryos, collected from *Tg(myl7:EGFP)^twu34^* fish, were injected in the yolk with 2 nL of the RNP working solution. Heart morphology was scored in the live embryos using the *myl7:EGFP* transgene. At 2 dpf, the hearts were scored for cardia bifida and other heart defects (e.g. incomplete looping, dysmorphic chambers), and at 4 dpf they were scored again to identify defects that had arisen since 2 dpf. The scoring data was analyzed using one-way ANOVA, comparing each experimental guide cocktail-injected group to controls, with Dunnett’s correction for multiple comparisons, and a cutoff of p < 0.05. Statistical analysis was performed and plots were generated in GraphPad Prism (version 10.0.3; Fig. 3D) or in MS Excel (Fig. 3C).

### Generation of zebrafish mutant strains for *pomp* and *psmd6*

To generate deletions encompassing the transcriptional start site and 5’ end of *pomp* and *psmd6*, we used a protocol based on that previously described [69]. CRISPR target sequences were selected using the Integrated DNA Technologies Alt-R HDR design tool (https://www.idtdna.com/site/order/designtool/index/HDRDESIGN). For each gene, 2 sites were chosen upstream and two downstream of the transcriptional start site, such that deletions of approximately 500-2500 base pairs would be generated. The Alt-R crRNAs were annealed with Alt-R tracrRNA to yield functional gRNA duplexes as in [69]. The four combinations of one upstream plus one downstream gRNA were tested to find the pair with the highest efficiency in generating the desired deletion. The sequences of the crRNAs used to generate the *pomp* and *psmd6* deletions are given in S8 Table. RNP complexes of gRNA + Cas9 protein (IDT Alt-R S.p. Cas9 Nuclease V3) were assembled as in [69], except that 0.5 µL each of one upstream and one downstream gRNA (each at 25 µM) were combined with 1 µL of 25 µM Cas9 plus 2 µL of H_2_O. After a 5-minute incubation at 37°C, 1 µL of phenol red injection solution was added. 1-cell stage embryos, collected from *Tg(myl7:EGFP)^twu34^* fish, were injected in the yolk with 1-2 nL doses of the RNP complexes. Injected embryos were raised to adulthood. To identify F0 fish with germline transmission of *pomp* or *psmd6* deletions, F0 adults were crossed with *Tg(myl7:EGFP)^twu34^* zebrafish. A subset of each of the resulting F1 clutches was screened by PCR analysis using primers flanking the expected deletions, where amplicons were only generated in animals carrying a deletion allele. The remaining F1 animals from positive F0 fish were raised to adulthood. Sanger sequencing was performed on the PCR-amplified deletion allele from F1 heterozygous animals and confirmed the deletion of the genomic sequence between the two CRISPR target sites. To identify heterozygous mutant carriers for the F1 and subsequent generations, fin-clippings from adults were collected, and PCR analysis was performed using a cocktail of three primers that generate different sized amplicons from the wild-type and deletion alleles. Primer sequences are provided in S8 Table. The deletion allele-specific primer pair spans the deletion but does not generate an amplicon from the wild-type allele due to the length of the intervening sequence, and the wild-type allele-specific primer pair includes one primer within the deletion. For *pomp^scm41^*, primers T1F1 + T1R1 produce a 225 bp product from the wild-type allele and T3F3 + T1R1 produce a 190 bp product from the deletion allele. For *psmd6^scm40^*, T3F1 + T3R1 produce a 298 bp product from the wild-type allele and T3F1 + T1R1 produce a 266 bp product from the deletion allele. The same PCR-based assays were used to genotype immunostained and RNA *in situ*-stained embryos. Heterozygous carriers were outcrossed with *Tg(myl7:EGFP)^twu34^* fish each generation.

### Quantitative reverse transcription PCR (qRT-PCR)

Embryos were obtained from incrosses of heterozygous *pomp^scm41^*or *psmd6^scm40^* fish. At 4 dpf, replicate groups of 10-20 phenotypically mutant and phenotypically wild-type larvae were flash-frozen in liquid nitrogen and then homogenized in TriZol (Invitrogen, ThermoFisher Scientific; 15596026) by trituration through a 27G needle. After phase separation, the aqueous portion was extracted with 24:1 chloroform:iso-amyl alcohol. One-half volume of 100% ethanol was added to the resulting aqueous phase and the samples were then further purified using the RNAqueous Micro Kit (Invitrogen, ThermoFisher Scientific; AM1931). Total RNA was reverse transcribed with the SensiFAST cDNA Synthesis kit (Bioline, Meridian Life Science; BIO-65054). Primer pairs were designed using Primer-BLAST such that they either span an intron (*pomp*) or one of the primers crosses an exon-exon boundary (*psmd6*). Primers for *rpl13a* were previously described [70]. Primers used are listed in S8 Table. qPCR was performed using the KAPA SYBR FAST kit (Roche; 07959567001) on a Bio-Rad CFX96 machine. Ct values for the genes of interest were normalized to *rpl13a,* and then ΔΔCts were calculated by normalizing each sample ΔCt to the average ΔCt of the replicate wild-type samples. The ΔΔCt values were log transformed to give fold change vs the average transcript level of the wild-type replicates and were plotted in GraphPad Prism 10. Un-paired, two-tailed *t*-tests were applied in GraphPad Prism 10 to compare wild-type and mutant samples.

### Bortezomib treatments

Embryos from spawnings of *Tg(myl7:EGFP)^twu34^* fish were enzymatically dechorionated with Pronase (Sigma; Cat. # 10165921001) at 24 hpf as described [67]. The embryos were arrayed in the wells of a 12-well plate at 25 animals per well in 3 mL of embryo medium (EM; 14.97 mM NaCl, 0.50 mM KCl, 0.98 mM CaCl_2_.2H_2_O, 0.15 mM KH_2_PO_4_, 0.99 mM MgSO_4_.7H2O, 0.05 mM Na_2_HPO_4_, 0.83 mM NaHCO_3_) containing bortezomib (Sigma; Cat. #5043140001) or 0.5% DMSO as a vehicle control. Treatments were done in triplicate wells. The drug treatments began at approximately 24 hpf, and the drug-containing media were replaced each day until 4 dpf, at which point the larvae were collected as 3 replicates per treatment condition and lysed as described below for immunoblotting.

### Immunoblotting

Embryos were obtained from incrosses of heterozygous *pomp^scm41^*or *psmd6^scm40^* fish. At 4 dpf, replicate groups of 20 phenotypically mutant and 20 phenotypically wild-type larvae were de-yolked in normal Ringer’s and lysed in 12 µL per embryo of 1.5x NuPAGE LDS sample buffer (Invitrogen, ThermoScientific; Cat. # NP0007). Approximately one embryo equivalent per sample was separated by reducing SDS-PAGE on 4-12% Bis-Tris NuPAGE gels (Invitrogen, ThermoScientific; Cat. # NP0322BOX) in MOPS buffer (Invitrogen, ThermoScientific; Cat. # NP0001) and transferred to nitrocellulose. Blots were blocked with Intercept TBS (LI-COR Biosciences; Cat. # 927-60001) and probed with anti-ubiquitinated proteins (clone FK2; 1:1000; EMD Millipore; Cat. # 04-263) followed by goat anti-mouse IR Dye 800CW secondary (1:10,000; LI-COR Biosciences; Cat. # 926-32210). Blots were imaged, then additionally probed with anti-alpha actin (CloneC4; 1:1000; MP Biomedical; Cat. # 0869100) followed by goat anti-mouse DyLight 680 secondary (1:20,000; ThermoScientific; Cat. # 35518) and were then re-imaged. Immunoblots were visualized on a LI-COR Odyssey infrared scanner, and images were generated using Image Studio Lite v5.2 (LI-COR Biosciences).

### Zebrafish whole-mount RNA in situ hybridization and immunostaining

The following cDNA probes were used for RNA in situ hybridization: *myl7* [44] and *elnb* [48]. Whole-mount in situ hybridization colorimetric and fluorescent in situ staining was performed as previously described [71,72], except that hybridizations for both experiments were performed in hybridization buffer that included 5% dextran sulfate. Following staining, tail clips from post-*in situ* hybridized embryos were lysed and genotyped.

The primary antibodies used for immunostaining were anti-α-actinin (ACTN2) (clone EA-53, 1:200; Sigma; Cat. # A7811) and anti-GFP (1:200; Torrey Pines Biolabs; Cat. # TP401). The secondary antibodies used were goat anti-rabbit IgG (H+L) Alexa Fluor 488 (Invitrogen, ThermoScientific; Cat. # A-11008) and goat anti-mouse IgG (H+L) Alexa Fluor 568 (Invitrogen, ThermoScientific; Cat. # A-11004). 96 hpf larvae were fixed in fresh 4% paraformaldehyde in PBS for 4 hours at room temperature, washed out of fixative, and stored in PBS containing 0.02% sodium azide at 4°C. The larvae were permeabilized with 5ug/mL Proteinase K (Sigma; Cat. # 3115836001) for 90 minutes, washed in PBS plus 0.1% Tween-20 (PBTw), treated with acetone for 20 minutes at –20°C, washed with PBTw, permeabilized with 1% sodium dodecyl sulfate in PBS for 15 minutes, and washed again with PBTw before being blocked in 2% heat-inactivated normal goat serum, 2% bovine serum albumin in PBTw (Fish Block) for 16 hours at 4°C. Primary antibodies were diluted in Fish Block and incubated with the embryos for 16-20 hours at 4°C. The embryos were then washed in PBTw, re-blocked in Fish Block for 2 hours at room temperature, and incubated with secondary antibodies plus 5µg/mL DAPI diluted in Fish Block for 16-20 hours at 4°C. Finally, the embryos were washed in PBTw and stored in 4% PFA at least overnight before tail tips were dissected for genotyping.

### Microscopic imaging of zebrafish embryos and larvae

Live embryos were anaesthetized in tricaine (Sigma; A5040) and then transferred to 2.5% methyl cellulose (Sigma; M0387) in ICS water. Whole-mount RNA colorimetric *in situs* were also imaged in 2.5% methyl cellulose in ICS water. In both cases, images were captured using an Olympus SZX16 stereomicroscope with an Olympus DP74 camera and cellSens Dimension v4.1 imaging software. Sequential brightfield and GFP fluorescence images were captured and later merged in Adobe Photoshop. For whole-mount immunostaining and fluorescent *in situs*, embryos were partially cleared in 80% glycerol. The trunk and tail were removed to facilitate mounting of the head and pharynx in 4% propyl gallate (Sigma; P3130) in 80% glycerol. Hearts were imaged on a Leica TCS SP5 confocal with a 20x air or 40x oil immersion objective.

Maximum intensity projections were made in ImageJ (https://fiji.sc/). Rounded cardiomyocytes were manually counted using the “Multi-point” tool in ImageJ by scrolling through Z-stacks captured with the 40x objective of merged GFP and DAPI channel images of individual hearts and marking the double-positive nuclei protruding outward from the myocardial wall. Cardiac sarcomere lengths and myofibril widths were measured using the “Straight line” tool and Measure function in ImageJ on the α-actinin channel of Z-stacks captured with the 40x objective. The measurements were made only where an individual myofiber could be clearly distinguished. Only one of each measurement was made per myofiber to ensure that a representative sample of myofibers per heart were scored. Rounded cell count data and sarcomere measurements were plotted in GraphPad Prism 10. Un-paired, two-tailed *t*-tests were applied in GraphPad Prism 10 to compare wild-type and mutant samples.

## Acknowledgements

We thank the SCRI Office of Animal Care for caring for the zebrafish. We thank Kylie Kerker for her efforts in the initial zebrafish CRISPR screening through the University of Washington School of Medicine Scholarship of Discovery program.

## Author Contributions

**Conceptualization:** DB, LM

**Data Curation:** GHFIII, WR, LM

**Formal Analysis:** GHFIII, WR, IY, MLL, DB, LM

**Funding Acquisition**: LM

**Investigation:** GHFIII, WR, IY, MLL, LM

**Methodology:** GHFIII, WR, LM

**Project Administration:** LM

**Resources:** LM

**Supervision**: LM

**Writing-Original Draft Preparation:** GHFIII, WR, LM

**Writing-Review and Editing:** GHFIII, WR, IY, MLL, DB, LM

## Supporting Information

**S1 Table. Known CHD gene list.**

**S2 Table. *s*_het_ top to deciles gene list.**

**S3 Table. Mouse embryonic heart-expressed gene list.**

**S4 Table. Candidate CHD gene list.**

**S5 Table. Candidate CHD genes with CHD *de novo* variants.**

**S6 Table. Candidate CHD genes with human embryonic heart expression.**

**S7 Table. Local network clusters.**

**S8 Table. Sequences of oligonucleotides used.**

